# HLA-MA: Simple yet powerful matching of samples using HLA typing results

**DOI:** 10.1101/066548

**Authors:** Clemens Messerschmidt, Manuel Holtgrewe, Dieter Beule

## Abstract

**Summary:** We propose the simple method HLA-MA for consistency checking in pipelines operating on human HTS data. The method is based on the HLA typing result of the state-of-the-art method Opti-Type. Provided that there is sufficient coverage of the HLA loci, comparing HLA types allows for simple, fast, and robust matching of samples from whole genome, exome, and RNA-seq data. This approach is reliable for sample re-identification even for samples with high mutational loads, e.g., caused by microsatellite instability or POLE1 defects.

**Availability and Implementation:** The software is implemented In Python 3 and freely available under the MIT license at https://github.com/bihealth/hlama and via Bioconda.

## 1 Introduction

The clinical applications of high-throughput sequencing (HTS) are diverse and manifold. While allowing for precise assays, HTS itself, of course, does not help with the more mundane problems in the life sciences such as sample swaps. While hard to detect in the general case, it is possible to use expected relations between two or more data sets for validation. In the case of analyzing data from pedigrees and matched tumor/normal pairs, relations are fairly simple. Matched tumor/normal pairs should have identical genotypes, or at least very similar ones as the rate of somatic mutation in tumors is expected to be much lower than the difference between two random samples from two different individuals. In pedigrees, Mendelian inheritance rules should hold true. One set of genes well suited for this task is the Human Leukocyte Antigen (HLA) system. The human HLA (class I) type consists of six alleles of the highly variable MHC class I genes. Overall, more than 11,000 different alleles are known today [1]. The combination of alleles will almost certainly be unique for any human individual and therefore the HLA type can be used as a fingerprint for any human sample and therefore even help to recover correct sample relationships.

## 2 Method

HLA-MA uses a simple, yet effective idea. For a list of HTS samples (each having single or paired-end, DNA-seq or RNA-seq data) and a description of their relation (e.g., being matched as tumor/normal pairs or their family relation in a pedigree), the HLA types are inferred. Then, they are compared up to two-digit (classical serotyping) and four-digit (protein sequence) precision with each other. Previous methods, e.g., as implemented by Samtools *gtcheck* [2] require variant calls from the samples. Such variant calls might be misleading in the case of high mutational load, e.g., in colon cancer data with microsatellite instability or POLE1 defects. In matched tumor/normal mode, a full match is expected (i.e., all HLA types should match), whereas in pedigree mode, the inheritance of the HLA genes is checked to follow the expected Mendelian inheritance rules.

HLA-MA is a Python 3 program that uses Snakemake [3] for the orchestration of its workflow. First, Yara [4] is used for prefiltering the reads and then OptiType [5] is used for performing the HLA typing, which internally uses RazerS3 [6] for read mapping. OptiType was chosen because it performed best on a selection of data sets compared to competing methods in our benchmarks (cf. S1 Supporting Information). The resulting HLA types are then simply compared in Python code. HLA-MA can also be run in parallel on a compute cluster (e.g., using Grid Engine) through the cluster parallelization support in Snakemake.

## 3 Results

For demonstration purposes, we obtained the FASTQ files from the ENA (European Nucleotide Archive) of the Illumina Platinum Genomes CEU pedigrees [7], ENA identifier ERP001960) and matched tumor/normal DNA and RNA-seq for cell line HCC1395 [8]. HLA-MA successfully confirmed the provided sample relation information. Table 1 shows typical running times of HLA-MA on different data set types. The running times were measured on a 16 core Xeon E5–2650 system (2 threads per core) where HLA-MA was run with 8 threads. Running time was dominated by the first Yara read alignment. All data sets were paired-end sequencing data with read lengths 2 × 101 bp.

**Table 1:** Data set properties, average running time and memory usage of the full HLA-MA pipeline

| data set | avg. read count per sample | avg. running time per sample | max. memory |
| --- | --- | --- | --- |
| HiSeq WGS | 2 x 800 M | 1h 15 min | 1.5 GB |
| Germline WES | 2 x 150 M | 15 min | 1.5 GB |
| Tumor WES | 2 x 300 M | 30 min | 1.5 GB |
| RNA-seq | 2 x 100 M | 15 min | 1.5 GB |

## 4 Discussion

We presented HLA-MA a simple and fast, yet effective method for the screening of raw Illumina HTS data for sample swaps for data sets where the HLA loci are covered. This is the case for a large range of clinical applications such as WES, WGS, and RNA-seq. Further, adding appropriate probes into specialized panels is usually possible at low cost, such that HLA-MA can also be used for such specialized applications. The HLA-MA pipeline takes a relatively short time and can be run before or in parallel with the usual downstream processing (on the same system used for benchmarking, the alignment step from FASTQ to sorted and indexed BAM takes about 14 h for genomes and 2-4 h for exomes with BWA-MEM). Thus, HLA-MA can be run together with the read alignment step without adding a considerable overhead and allowing for spotting tentative sample swaps early on in clinical pipelines. The computation of HLA types might even be practically for free if the research question requires HLA typing, as is the case in cancer immunology, for example.

## Supplementary Materials

S1 Supporting Information

